# Analysis of genetically driven alternative splicing identifies FBXO38 as a novel COPD susceptibility gene

**DOI:** 10.1101/549626

**Authors:** Aabida Saferali’, Jeong H. Yun, Margaret M. Parker, Phuwanat Sakornsakolpat, Robert P. Chase, Andrew Lamb, Brian D. Hobbs, Marike H. Boezen, Xiangpeng Dai, Kim de Jong, Terri H. Beaty, Wenyi Wei, Xiaobo Zhou, Edwin K. Silverman, Michael H. Cho, Peter J. Castaldi, Craig P. Hersh, Investigators COP DG ene Investigators, and the International COPD Genetics Consortium

## Abstract

While many disease-associated single nucleotide polymorphisms (SNPs) are associated with gene expression (expression quantitative trait loci, eQTLs), a large proportion of complex disease genome-wide association study (GWAS) variants are of unknown function. Some of these SNPs may contribute to disease by regulating gene splicing. Here, we investigate whether SNPs that are associated with alternative splicing (splice QTL or sQTL) can identify novel functions for existing GWAS variants or suggest new associated variants in chronic obstructive pulmonary disease (COPD).

RNA sequencing was performed on whole blood from 376 subjects from the COPDGene Study. Using linear models, we identified 561,060 unique sQTL SNPs associated with 30,333 splice sites corresponding to 6,419 unique genes. Similarly, 708,928 unique eQTL SNPs involving 15,913 genes were detected at 10% FDR. While there is overlap between sQTLs and eQTLs, 60% of sQTLs are not eQTLs. Co-localization analysis revealed that 7 out of 21 loci associated with COPD (p<1×10^−6^) in a published GWAS have at least one shared causal variant between the GWAS and sQTL studies. Among the genes identified to have splice sites associated with top GWAS SNPs was *FBXO38*, in which a novel exon was discovered to be protective against COPD. Importantly, the sQTL in this locus was validated by qPCR in both blood and lung tissue, demonstrating that splice variants relevant to lung tissue can be identified in blood. Other identified genes included *CDK11A* and *SULT1A2*.

Overall, these data indicate that analysis of alternative splicing can provide novel insights into disease mechanisms. In particular, we demonstrated that SNPs in a known COPD GWAS locus on chromosome 5q32 influence alternative splicing in the gene *FBXO38*.

**Author Summary:** While it is known that chronic obstructive pulmonary disease (COPD) is caused in part by genetic factors, few studies have identified specific causative genes. Genetic variants that alter the expression levels of genes have explained part of the genetic component of COPD, however, there are additional genetic variants with unknown function. In some genes the protein coding sequence can be altered by a mechanism known as RNA splicing. We hypothesized that some genetic variants that are associated with risk of COPD contribute to the disease by altering RNA splicing. In this study, we identified genetic variants that are associated both with COPD risk and RNA splicing. In particular, we found that a COPD associated variant of previously unknown function may contribute to the inclusion of a new exon in the *FBXO38* gene. These finding are significant because they indicate that analysis of RNA splicing can help identify genes that contribute to disease.

## Introduction

Chronic obstructive pulmonary disease (COPD) is characterized by irreversible airflow obstruction. While cigarette smoking is the leading environmental risk factor for COPD, only a subset of smokers develop the disease. Genetic factors have been shown to contribute to COPD susceptibility, with the best characterized example being *SERPINA1*, the causal gene for α1-antitrypsin deficiency (1–5).

Genome-wide association studies (GWAS) have been used to identify common genetic variants associated with many complex diseases, including COPD. These studies have identified multiple well-replicated genome-wide significant loci, including a locus on 15q25 (*CHRNA3/CHRNA5/IREB2*), *FAM13A*, *HHIP*, *CYP2A6* and *HTR4* (6–10). In addition, a recent large meta-analysis identified twenty-two genome-wide significant loci in COPD, of which 13 were newly genome-wide significant (11). However, the majority of single nucleotide polymorphisms (SNPs) that have been identified in COPD GWAS are of unknown function. Since most GWAS variants are located in noncoding regions, it is likely that these variants contribute to COPD susceptibility through transcriptional regulation of target genes.

Expression quantitative trait locus (eQTL) studies have been used to identify SNPs that contribute to gene expression levels, thereby validating GWAS associations in addition to providing insight into the biological mechanism responsible. To date, only a moderate percentage of GWAS findings have been shown to be eQTLs with strong effect size (12). This is likely due to a variety of mechanisms including inadequate sample size, multiple testing correction, tissue type investigated, as well as the focus on total gene-level expression levels without consideration of transcript isoforms.

COPD has been shown to have a disproportionate amount of alternative splicing compared to other complex diseases such as type 2 diabetes, Alzheimer’s disease, and Parkinson’s disease, suggesting that transcriptional regulation through splicing may play an important role (13). Genetic variation could contribute by altering mRNA splicing, which in turn could result in changes in protein sequence or expression levels. Several recent studies have demonstrated that SNPs associated with alternative splicing (sQTLs) are enriched for GWAS variants. Evidence suggests that at least 20-30% of disease causing mutations may affect pre-mRNA splicing (14, 15). Furthermore, several reports have discovered that there are a proportion of GWAS SNPs that have evidence of sQTLs but not eQTLs, indicating that sQTL analysis can provide additional insight into the functional mechanisms underlying GWAS results (16) (17, 18).

Here, we characterize sQTLs in human peripheral blood in COPD to determine whether these loci can identify novel functions for COPD GWAS variants. We hypothesize that a substantial fraction of COPD GWAS loci influence disease susceptibility through sQTLs. Previous studies to identify sQTLs have been performed in small sample sizes or have focused on exon expression instead of differential exon usage (16). This study characterizes variants associated with differential exon usage in a large population.

## Materials and Methods

### Study Population

This study included 376 non-Hispanic white subjects from the COPDGene study. COPDGene enrolled individuals between the ages of 45 and 80 years with a minimum of 10 pack-years of lifetime smoking history from 21 centers across the United States (19). These subjects returned for a second study visit 5 years after the initial visit at which time they completed additional questionnaires, pre-and post-bronchodilator spirometry, computed tomography of the chest, and provided blood for complete blood counts (CBCs) and RNA sequencing. In this study, moderate to severe COPD was defined as GOLD spirometric grades 2-4 (20).

### RNASeq data acquisition and processing

The protocol for RNASeq data generation has been previously described (21). Total RNA was extracted from PAXgene Blood RNA tubes using the Qiagen PreAnalytiX PAXgene Blood miRNA Kit (Qiagen, Valencia, CA). Extracted samples with a RIN > 7 and concentration > =25ug/uL were included in sequencing. Globin reduction, ribosomal RNA depletion and cDNA library prep was performed using the TruSeq Stranded Total RNA with Ribo-Zero Globin kit (Illumina, Inc., San Diego, CA). The Illumina HiSeq 2500 was used to generate 75 bp reads, and an average of 20 million reads were generated per sample.

### Read alignment, mappability filtering and quality control

Reads were trimmed using skewer (22) to remove specified TruSeq adapter sequences. Quality control was performed using the FASTQC (23) and RNA-SeQC (24) programs. Trimmed reads were aligned to the GRCh37 reference genome using a two-pass alignment method with STAR 2.5 (25). Following sequence alignment, mappability filtering to correct for allelic bias in read mapping was performed using WASP (26). An average of 840,000 reads were removed due to read mapping bias resulting in an average of 19.9 million reads being available for subsequent analysis.

### Genotyping

Genotyping was performed by Illumina (San Diego, CA) on the HumanOmniExpress Array. Eagle v. 2.3 was used for phasing and HRC reference panel version 1.1 was used for imputation. SNPs with minor allele frequency greater than 0.05 and imputation R^2^ greater than 0.5 were included in analysis (9, 11).

### Quantification of Splicing Ratios and Gene Expression Counts

Gene expression counts were computed using Rsubread (27). Quantification of splicing ratios was performed using Leafcutter (17). This method extracts junctional reads (or reads that span introns) from aligned bam files and clusters them according to shared start or stop positions. Default leafcutter parameters were used in the detection of clusters, i.e., 50 split reads across all individuals were required to support each cluster, and introns up to 500 kb were included. For sQTL analysis, intron ratios were calculated by determining how many reads support a given exon-intron junction in relation to the number of reads in that region. Introns used in less than 40% of individuals were filtered out, and the remaining intron ratios were used as input for sQTL analysis.

### eQTL and sQTL Analysis

MatrixeQTL (28) was used to test for association between genotype of all SNPs within 1000 kb of a gene (cis-) and quantifications of gene expression or alternative splicing using linear models, adjusting for age, gender, pack-years of smoking, current smoking status, white blood cell differential, PEER factors of expression data and principal components of genetic ancestry. A total of 5,405,234 SNPs were tested for association with 25,313 genes and 97,365 splice sites.

### SNP lookup and colocalization analysis

COPD-associated SNPs were obtained from a subset of a published GWAS (11). We selected 920 SNPs with p<1 × 10^−6^ in white subjects to match the ethnicity of the RNA-Seq data. These SNPs were grouped into 21 loci based on their genomic positions. SNPs that were associated with splicing at the 10% FDR, or with gene expression at the 10% FDR were identified. The 10% FDR threshold was selected based on published sQTL studies (18, 29). Co-localization analysis was performed for GWAS loci that contained sQTLs using eCAVIAR (30).

### Quantitative PCR (qPCR) of *FBXO38*

Thirty COPDGene blood samples were selected for qPCR based on expression levels in RNASeq data. An additional ninety resected lung tissue samples from individuals undergoing thoracic surgery (31) were selected based on genotype. A total of 400 ng of RNA was reverse transcribed using the SuperScript III First-Strand Synthesis System (ThermoFisher Scientific, Waltham, MA) for blood RNA or the iScript™ cDNA Synthesis Kit (BioRad, Hercules, CA) for lung RNA. A Taqman assay was designed to amplify cDNA fragments from the 3’ region of the cryptic *FBX038* exon, to the 5’ region of exon 10 (Supplementary Figure 1). A predesigned Taqman assay (Hs01004563_mH) was used to amplify the alternate isoforms from the 3’ region of exon 9 to the 5’ region of exon 10. A final amount of 30 ng of cDNA was amplified per well, and each sample was assayed in triplicate. *GAPDH* was amplified as a housekeeping gene to control for RNA concentration. To calculate the ratio of transcripts containing the novel exon, delta CT values for the novel isoform were divided by delta CT values for the alternate isoform. Linear regression was performed to test for an additive relationship between genotype and splicing ratio.

### Immunoblots and Immunoprecipitation

Cells were lysed in EBC buffer (50 mMTris pH 7.5, 120 mMNaCl, 0.5% NP-40) supplemented with protease inhibitors (Complete Mini, Roche) and phosphatase inhibitors (phosphatase inhibitor cocktail set I and II, Calbiochem). The protein concentrations of lysates were measured by the Beckman Coulter DU-800 spectrophotometer using the Bio-Rad protein assay reagent. Same amounts of whole cell lysates were resolved by SDS-PAGE and immunoblotted with indicated antibodies. For immunoprecipitation, 1000 μg whole cell lysates were incubated with the indicated anti-Flag M2 affinity gel and monoclonal anti-HA agarose for 3-4 hr at 4 degree (Millipore Sigma). Immunoprecipitants were washed five times with NETN buffer (20 mMTris, pH 8.0, 100 mMNaCl, 1 mM EDTA and 0.5% NP-40) before being resolved by SDS-PAGE and immunoblotted with indicated antibodies.

## Results

### Identification of sQTLs and eQTLs in human peripheral blood

This analysis included 376 Non-Hispanic white COPDGene participants (Supplementary Table 1). Samples were sequenced to a depth of approximately 20 million reads per sample, and splice ratios from a total of 97,365 splice clusters (defined as overlapping introns that share a spice donor or acceptor site) were included in the sQTL analysis. Gene expression counts for 25,312 genes were used for eQTL detection.

We identified 1,706,704 cis-sQTLs at 10% FDR, comprising 561,060 unique SNPs (Supplementary Table 2). These SNPs were associated with 30,333 splice sites which were annotated to 6742 unique genes (Supplementary Table 3). Similarly, we identified 1,242,993 cis-eQTLs corresponding to 708,928 unique SNPs. These SNPs were associated with expression of 15,913 genes. We found that 44.6% of sQTLs were also eQTLs, but that 55.3% (310,361 SNPs) were sQTLs exclusively. In addition, 2299 genes contained at least one splice site that was significantly associated with genotype of a neighboring SNP, while total gene expression of the same gene was not significantly associated with any SNP. This suggests that analysis of sQTLs can identify novel regulatory events that are not captured through whole gene expression analysis.

### Pathway analysis of genes with sQTLs

To characterize the biological functions of genes for which alternative splicing was associated with nearby SNPs, Sigora (32) was used to identify overrepresented pathways in genes that had sQTLs but not eQLTs. This method of pathway analysis focuses on genes or gene pairs that are specific to a single pathway. In this way it utilizes the status of other genes in the experimental context to identify the most relevant pathways and minimize the identification of spurious pathways. To minimize the number of input genes, a conservative FDR cutoff of 5% was applied in the identification of genes with sQTLs. We identified 1752 genes which had significant sQTLs but not eQTLs at the 5% FDR. These genes were enriched for 20 KEGG pathways and 33 Reactome pathways (Supplementary Table 5 and 6), mostly related to RNA processing.

### Functional categories of cis eQTLs and sQTLs

Both eQTLs and sQTLs were categorized on the basis of their location relative to the gene with which they were associated, and the results are shown in Table 1. The genomic distribution of sQTLs and eQTLs was similar, with the majority located in intergenic (38% of sQTLs and 41% of eQTLs) and intronic (50% of sQTLs and 46% of eQTLs) regions. Only a small number of sQTLs (n=68) and eQTLs (n=74) were located in splice sites, defined as within an intron and 2 bp of an exon/intron boundary. In addition, we identified 7 sQTLs and 3 eQTLs located within an exon and 2 bp of an exon/intron boundary.

**Table 1:**
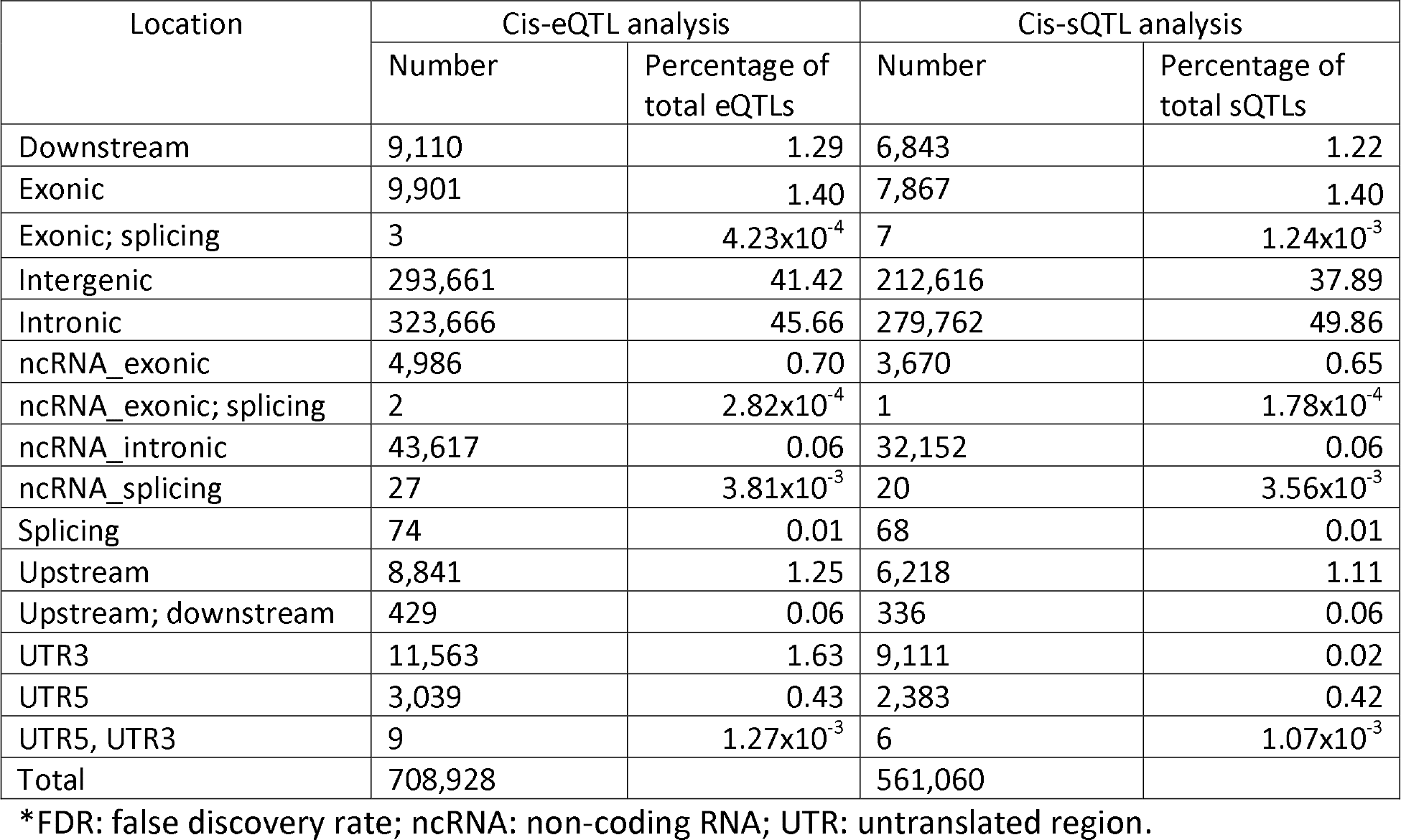
Functional annotation of cis expression quantitative trait loci (eQTLs) and splice QTLs (10% FDR), based on RefSeq annotation

| Location | Cis-eQTL analysis |  | Cis-sQTL analysis |  |
| --- | --- | --- | --- | --- |
|  | Number | Percentage of total eQTLs | Number | Percentage of total sQTLs |
| Downstream | 9,110 | 1.29 | 6,843 | 1.22 |
| Exonic | 9,901 | 1.40 | 7,867 | 1.40 |
| Exonic; splicing | 3 | $4.23 \times 10^{-4}$ | 7 | $1.24 \times 10^{-3}$ |
| Intergenic | 293,661 | 41.42 | 212,616 | 37.89 |
| Intronic | 323,666 | 45.66 | 279,762 | 49.86 |
| ncRNA_exonic | 4,986 | 0.70 | 3,670 | 0.65 |
| ncRNA_exonic; splicing | 2 | $2.82 \times 10^{-4}$ | 1 | $1.78 \times 10^{-4}$ |
| ncRNA_intronic | 43,617 | 0.06 | 32,152 | 0.06 |
| ncRNA_splicing | 27 | $3.81 \times 10^{-3}$ | 20 | $3.56 \times 10^{-3}$ |
| Splicing | 74 | 0.01 | 68 | 0.01 |
| Upstream | 8,841 | 1.25 | 6,218 | 1.11 |
| Upstream; downstream | 429 | 0.06 | 336 | 0.06 |
| UTR3 | 11,563 | 1.63 | 9,111 | 0.02 |
| UTR5 | 3,039 | 0.43 | 2,383 | 0.42 |
| UTR5, UTR3 | 9 | $1.27 \times 10^{-3}$ | 6 | $1.07 \times 10^{-3}$ |
| Total | 708,928 |  | 561,060 |  |

\*FDR: false discovery rate; ncRNA: non-coding RNA; UTR: untranslated region.

### Identification of genetic loci containing both GWAS associated SNPs and sQTLs

sQTLs were at least as enriched among low p-value associations with COPD case status as eQTLs (Supplementary Figure 2). To identify whether sQTLs can identify functions for GWAS SNPs that could not be explained by eQTLs alone, 920 SNPs associated with COPD in white subjects with a p <1 × 10^−6^ were grouped into 21 genetic loci according to genomic position (Table 2). These SNPs were then interrogated in the eQTL and sQTL data sets to identify which GWAS SNPs were also associated with alternative splicing or gene expression at 10% FDR. Of these 920 SNPs, 67 SNPs were both sQTLs and eQTLs, 71 SNPs were eQTLs alone, and 156 were sQTLs alone, indicating that a greater number of GWAS SNPs are sQTLs than eQTLs. Out of the 21 genomic loci, 6 included GWAS-associated SNPs that were eQTLs, and 7 included GWAS SNPs that were sQTLs (Table 2). There were three loci that contained sQTLs but not eQTLs for any gene

**Table 2:**
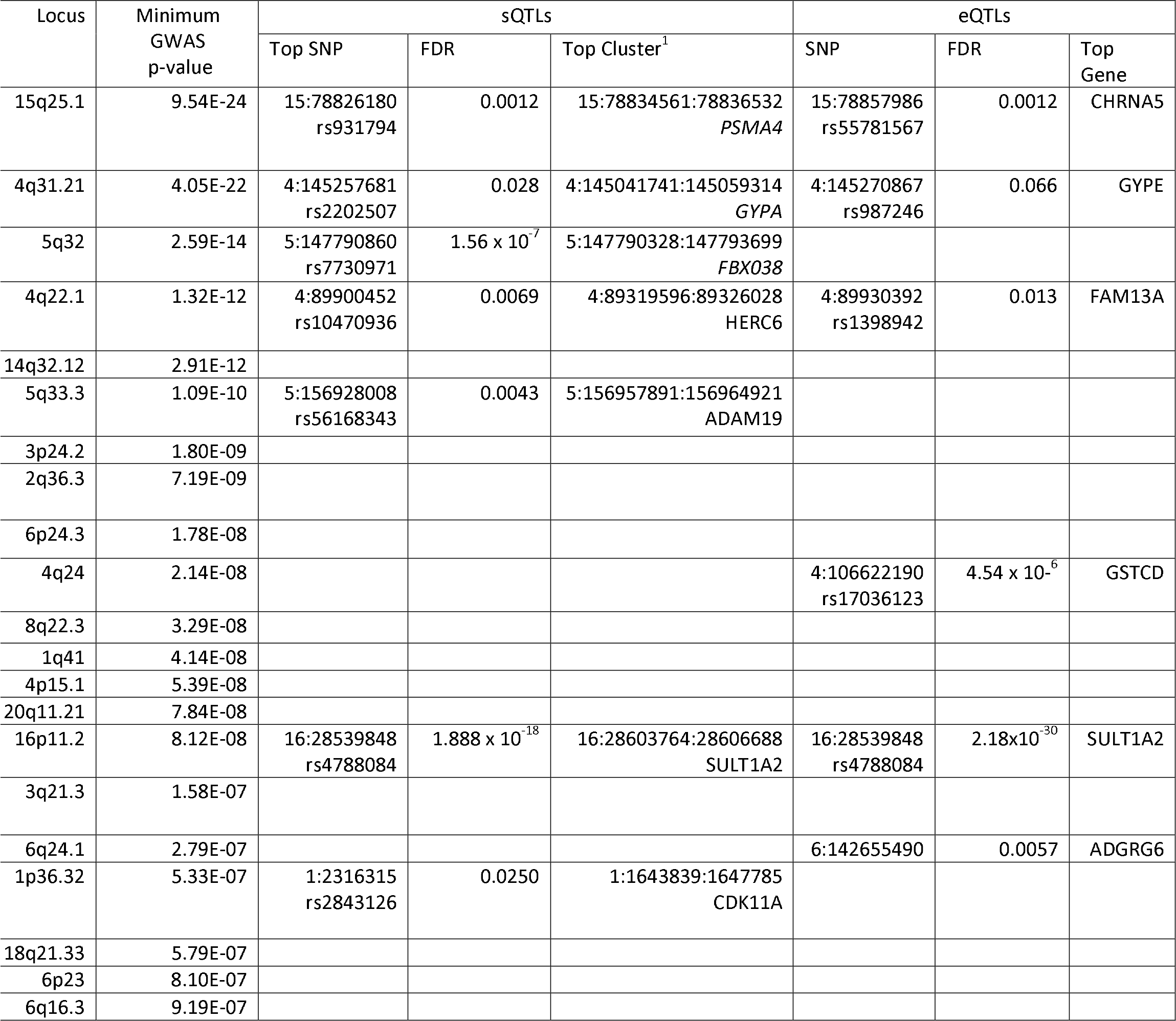
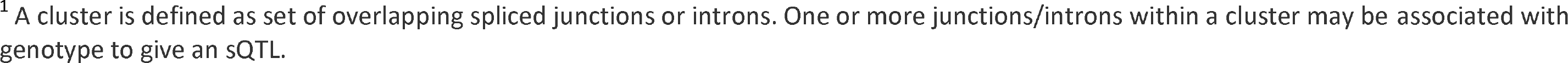
COPD genome-wide association study loci containing cis-eQTLs and cis-sQTLs at 10% FDR

| Locus | Minimum GWAS p-value | sQTLs |  |  | eQTLs |  |  |
| --- | --- | --- | --- | --- | --- | --- | --- |
|  |  | Top SNP | FDR | Top Cluster<sup>1</sup> | SNP | FDR | Top Gene |
| 15q25.1 | 9.54E-24 | 15:78826180<br>rs931794 | 0.0012 | 15:78834561:78836532<br>*PSMA4* | 15:78857986<br>rs55781567 | 0.0012 | CHRNA5 |
| 4q31.21 | 4.05E-22 | 4:145257681<br>rs2202507 | 0.028 | 4:145041741:145059314<br>*GYPA* | 4:145270867<br>rs987246 | 0.066 | GYPE |
| 5q32 | 2.59E-14 | 5:147790860<br>rs7730971 | $1.56 \times 10^{-7}$ | 5:147790328:147793699<br>*FBX038* | | | |
| 4q22.1 | 1.32E-12 | 4:89900452<br>rs10470936 | 0.0069 | 4:89319596:89326028<br>*HERC6* | 4:89930392<br>rs1398942 | 0.013 | FAM13A |
| 14q32.12 | 2.91E-12 |  |  |  |  |  |  |
| 5q33.3 | 1.09E-10 | 5:156928008<br>rs56168343 | 0.0043 | 5:156957891:156964921<br>*ADAM19* |  |  |  |
| 3p24.2 | 1.80E-09 |  |  |  |  |  |  |
| 2q36.3 | 7.19E-09 |  |  |  |  |  |  |
| 6p24.3 | 1.78E-08 |  |  |  |  |  |  |
| 4q24 | 2.14E-08 | | | | 4:106622190<br>rs17036123 | $4.54 \times 10^{-6}$ | GSTCD |
| 8q22.3 | 3.29E-08 |  |  |  |  |  |  |
| 1q41 | 4.14E-08 |  |  |  |  |  |  |
| 4p15.1 | 5.39E-08 |  |  |  |  |  |  |
| 20q11.21 | 7.84E-08 |  |  |  |  |  |  |
| 16p11.2 | 8.12E-08 | 16:28539848<br>rs4788084 | $1.888 \times 10^{-18}$ | 16:28603764:28606688<br>*SULT1A2* | 16:28539848<br>rs4788084 | $2.18 \times 10^{-30}$ | SULT1A2 |
| 3q21.3 | 1.58E-07 |  |  |  |  |  |  |
| 6q24.1 | 2.79E-07 |  |  |  | 6:142655490 | 0.0057 | ADGRG6 |
| 1p36.32 | 5.33E-07 | 1:2316315<br>rs2843126 | 0.0250 | 1:1643839:1647785<br>*CDK11A* |  |  |  |
| 18q21.33 | 5.79E-07 |  |  |  |  |  |  |
| 6p23 | 8.10E-07 |  |  |  |  |  |  |
| 6q16.3 | 9.19E-07 |  |  |  |  |  |  |

<sup>1</sup> A cluster is defined as set of overlapping spliced junctions or introns. One or more junctions/introns within a cluster may be associated with genotype to give an sQTL.

### Colocalization analysis of SNPs in HTR4/FBXO38

To investigate whether GWAS associations may be attributed to alternative splicing, colocalization analysis was performed using eCAVIAR in the seven genetic loci containing both GWAS SNPs and sQTLs (Table 3). All seven of the GWAS loci had at least one variant with significant colocalization to sQTL data (colocalization posterior probability [CLPP] > 0.01) (Table 3 and Supplementary Table 4). The locus with the strongest colocalization between either GWAS & sQTL or GWAS & eQTL was located in the 5q32 region containing *HTR4/FBXO38*. This region was identified in the COPD case control GWAS, with 151 SNPs with p-value < 1×10^−6^; rs3995091 SNP located in the *HTR4* gene had the lowest p-value (2.59 × 10^−14^). Additional significant variants from an independent association in this region include rs7730971 and rs4597955 (Figure 1a) with GWAS p-values of 5.51×10^−12^ and 4.78 × 10^−12^, respectively. rs7730971 was significantly associated with splice sites in the *FBXO38* gene (Supplementary Figure 3), while this SNP was not associated with total expression of any gene. eCAVIAR analysis revealed that rs7730971 colocalized with the sQTL and GWAS data (CLPP = 0.845) (Figure 1b), while there was no colocalization with eQTLs for *FBXO38*, suggesting that the GWAS association may be caused be alternative splicing. Furthermore, eCAVIAR identified rs7730971 to be the SNP with the highest degree of colocalization between sQTL and GWAS data, suggesting that this may be the causative variant (Table 3 and Supplementary Table 4). Despite being the GWAS SNP with the minimal p-value in the locus, there was no colocalization between sQTL and GWAS data for rs3995091 (CLPP=0.002). Characterization of the splicing cluster associated with genotype of rs7730971 revealed a previously unannotated cryptic exon located at chromosome 16: 147,790,643 - 147,790,801. This 158 bp exon is present in a greater proportion of subjects with the GG genotype (13%) than the CC genotype (8%) (Figure 1c). The CC genotype is also associated with greater risk of COPD, so the novel isoform may be protective against COPD. Quantitative PCR independently validated the existence of the novel exon as well as replicated the effect of genotype on exon inclusion levels in RNA from whole blood(p=0.01, Figure 1d). Furthermore, the novel exon was identified in RNA from homogenized lung tissue, where splicing levels were also associated with rs7730971 genotype (p=0.007)(Figure 1d). The cryptic exon leads to a premature stop codon in *FBXO38*, which could alter or inhibit protein function. Additionally, immunoprecipitation was performed in 293T cells which indicated that FBXO38, an F-box protein with unknown substrates, interacts with Cullin 1, but not other Cullin members (Supplementary Figure 4), indicating that it may be a component of a SKP1-Cullin-1-F-box (SCF) type of E3 ubiquitin ligase complex. In combination, these findings suggest that the GWAS association at 5q32 may be partly explained by the inclusion of an exon which results in a truncated FBXO38 protein.

**Table 3:**
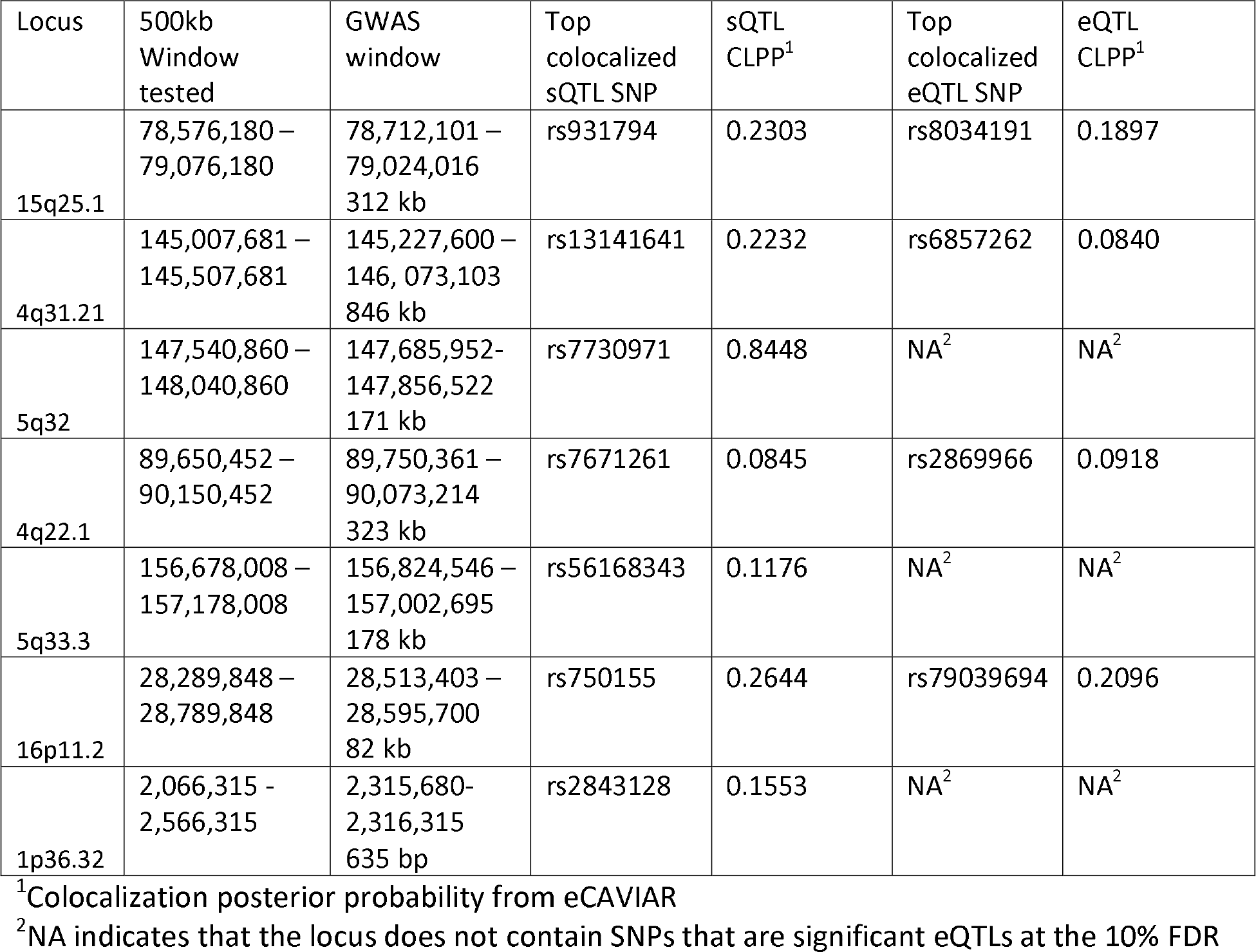
Colocalization of sQTLs and eQTLs with COPD case-control GWAS data

**Figure 1:**
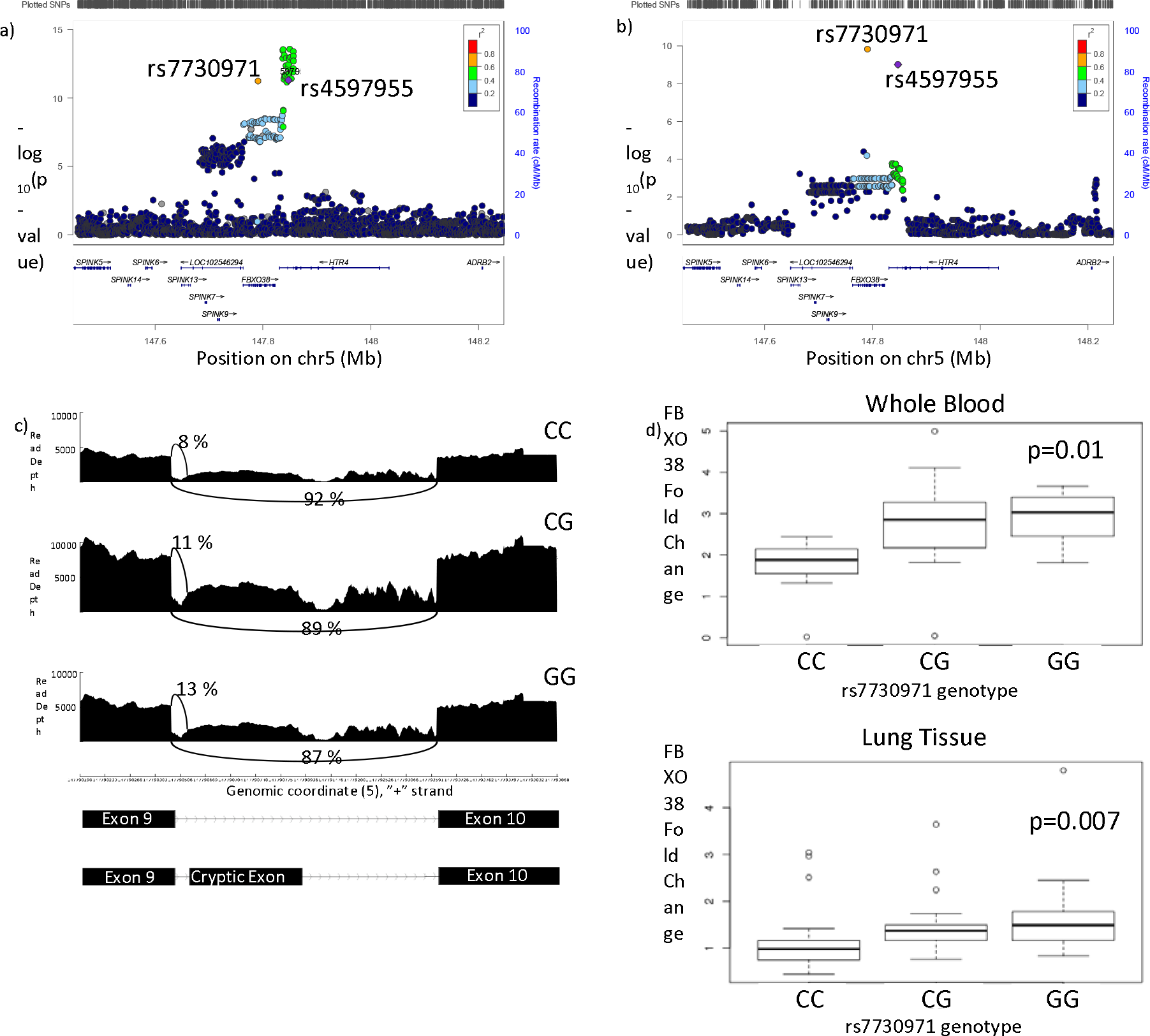
Colocalization analysis of sQTLs and COPD GWAS data at 5q32. a) Locus zoom plot of the GWAS association at 5q32. The secondary GWAS association at this locus consists of two SNPs – rs7730971 and rs4597955. The primary association is in moderate LD (R^2^ = 0.4-0.6) with this association. b) Locus zoom plot of sQTL data for the association between *FBXO38* splicing with genotype. SNPs associated with FBXO38 splicing are located in *HTR4* and *FBXO38* c) Visualization of the FBXO38 splice site associated with rs7730971 genotype. The y axis shows read depth and the x axis shows genomic position on chromosome 5. Arched lines indicate junction spanning reads. There are fewer junctional reads supporting the presence of the cryptic exon in the CC genotype (8%) compared to the GG genotype (13%) d) Boxplot of qPCR results showing the fold change of the isoform containg the crypic exon compared to the CC genotype in whole blood (n=30; selected based on expression levels) and lung tissue (n=90, selected based on genotype). The center line indicates median, the box captures the interquartile range, and the whiskers show the range excluding outliers.

### Colocalization analyses at additional GWAS loci

#### 16p11.2 – SNPs in CCDC101 are associated with both splicing and gene expression of SULT1A2

An 82 kb region on chromosome 16p11.2 was significantly associated with COPD in the GWAS (5 SNPs with p<1 × 10^−6^, Figure 2a). The rs4788084 SNP, which is in high linkage disequilibrium (LD) with the top GWAS SNP (rs7186573, R^2^ = 0.842), was significantly associated with both splicing and gene expression of *SULT1A2* (Supplementary Figure 5). The GWAS data colocalized with both sQTL and eQTL data (Figure 2b, Table 3, Supplementary Table 4), suggesting that the causal gene for the GWAS association is *SULT1A2* and that both gene expression and splicing may contribute to disease. The SNP with the greatest colocalization between GWAS data and sQTL data was rs750155 (CLPP=0.26), but for the eQTL data the greatest colocalization was with rs79039694 (CLPP=0.21) (Table 3). Characterization of the splicing cluster associated with genotype demonstrated that Exon 4 is differentially included based on rs4788084 genotype (Figure 2c). 81% of *SULT1A2* transcripts in individuals with the CC genotype included this exon, while it was present in 98% of transcripts in TT individuals. Exon 4 is skipped in one known transcript, *SULT1A2-002*. The T allele, which is associated with greater inclusion of exon 4, is associated with greater risk of COPD (based on GWAS data); increased expression of *SULT1A2-002* may be protective due to skipping of exon 4.

**Figure 2:**
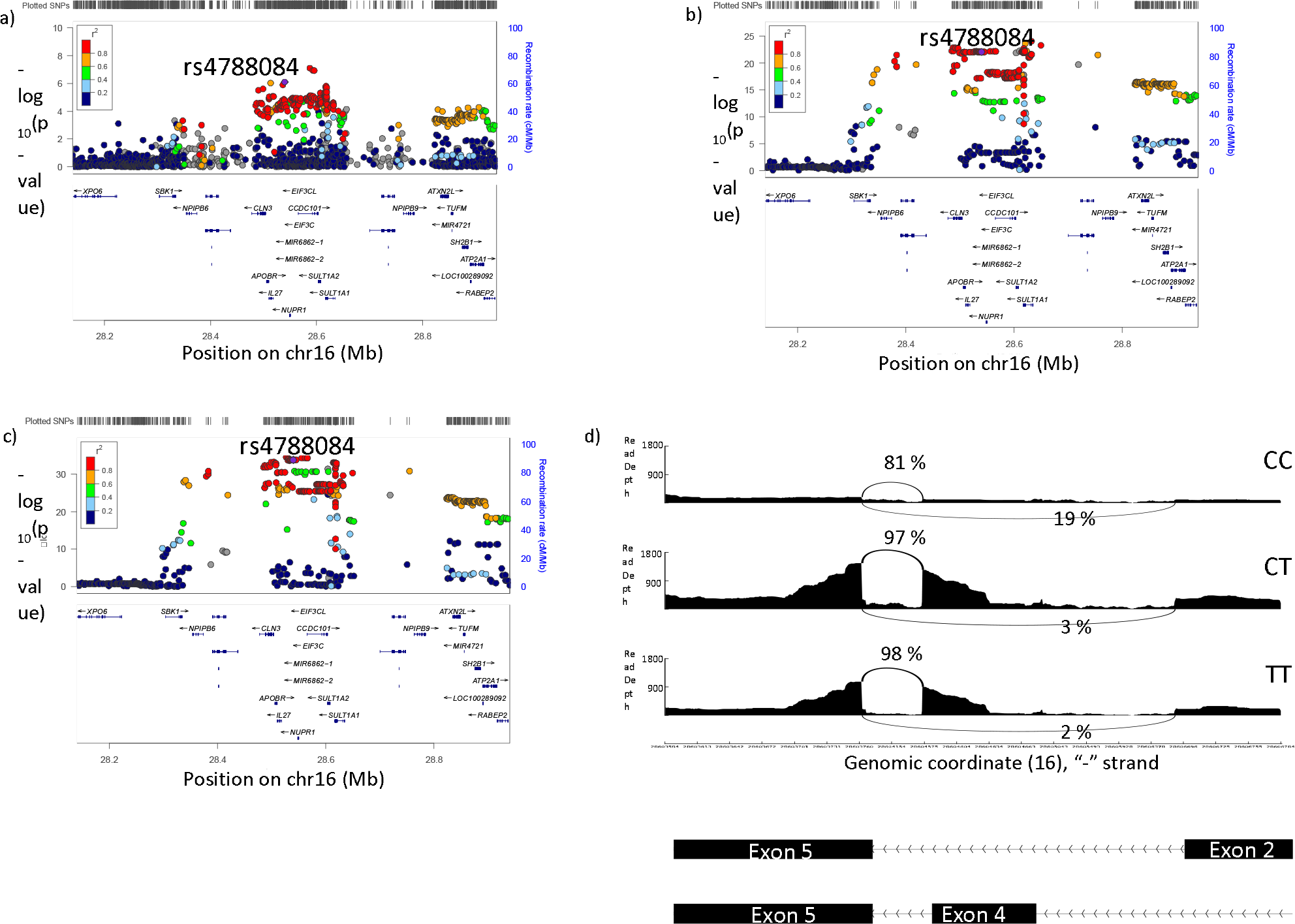
Colocalization analysis of sQTLs and eQTLs with GWAS data at 16p11.2. a) Locus zoom plot of the GWAS association at 16p11.2. b) Locus zoom plot of sQTL data for the association between *SULT1A2* splicing with genotype c) Locus zoom plot of eQTL data for the association between *SULT1A2* whole gene expression with genotype d) Visualization of the *SULT1A2* splice site associated with rs4788084 genotype. The y axis shows read depth and the x axis shows genomic position on chromosome 16. Arched lines indicate junction spanning reads.

#### 1p36.32 – SNPs in MORN1 are associated with splicing of CDK11A but not with gene expression

The COPD GWAS identified two SNPs in a 635 bp region at 1p36.32 that were associated at near-genome wide significance in whites (p = 5.33 × 10^−7^ and 8.24 × 10^−7^, Figure 3a). These SNPs are located in the *MORN1* gene, but are significantly associated with a splice site in *CDK11A* (Supplementary Figure 6). These SNPs were not associated with expression of any gene. Both GWAS SNPs colocalized with sQTL data (CLPP for rs2843128=0.155, rs2843126=0.120) (Figure 3b, Table 3, Supplementary Table 4). This indicates that splicing of *CDK11A* may be causative of the GWAS association. The identified splice site corresponds to an annotated exon in *CDK11A* (Figure 3c). This exon is skipped in one known transcript – *CDK11A-201*. The minor allele of rs2843126 (A) is associated with greater levels of inclusion of exon 6, with 15% of *CDK11A* transcripts including this exon in AA individuals, vs 12% inclusion rates in GG individuals. Furthermore, the A allele is associated with greater risk of COPD. This suggests that increased expression of the CDK11A-201, which skips exon 6, may be protective against COPD.

**Figure 3:**
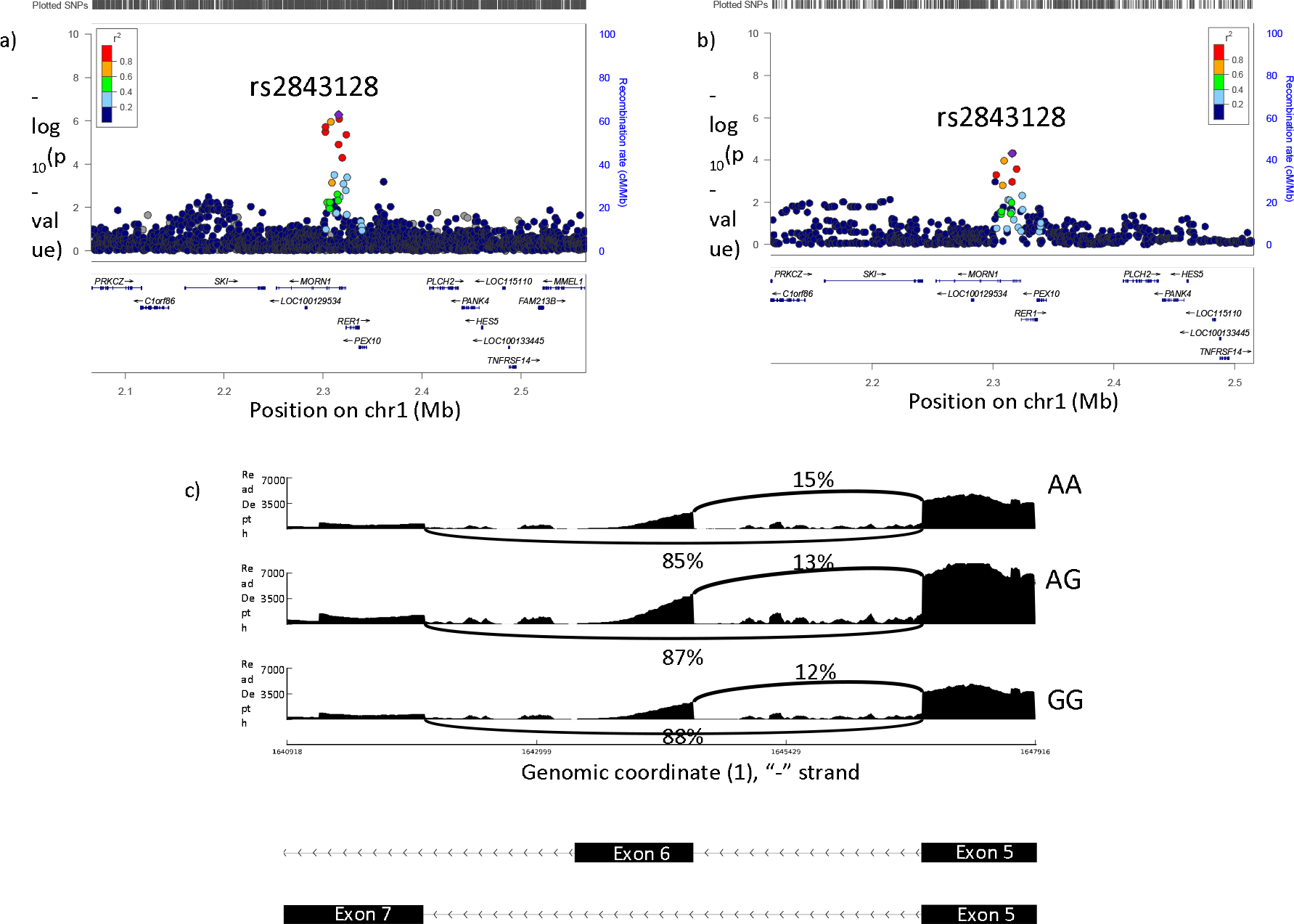
Colocalization analysis of sQTLs and COPD GWAS data at 1p36.32. a) Locus zoom plot of the GWAS association at 1p36.32. b) Locus zoom plot of sQTL data for the association between *CDK11A* splicing with genotype c) Visualization of the *CDK11A* splice site associated with rs2843128. The y axis shows read depth and the x axis shows genomic position on chromosome 1. Arched lines indicate junction spanning reads.

## Discussion

eQTL studies can provide insight into the biological mechanisms responsible for disease associations. While many disease-associated SNPs are eQTLs, a large proportion of GWAS variants are of unknown function. SNPs that are associated with transcript isoform variation may contribute to disease risk and explain additional GWAS associations. In this study, we characterized eQTLs and sQTLs in human peripheral blood to identify novel functions for COPD GWAS associations. Among the genes identified to have splice sites associated with GWAS SNPs were *CDK11A*, which may be of biological relevance for COPD due to its role in apoptosis; *SULT1A2* which has both eQTLs and sQTLs which strongly colocalize to the GWAS signal; and of particular interest, *FBXO38*, in which a novel exon is protective against COPD.

Although we found a larger number of eQTLs than sQTLs genome wide (708,928 vs 561,060 unique SNPs), a greater number of COPD GWAS SNPs were sQTLs (156/920) than eQTLs (89/920) This phenomenon has previously been shown in multiple sclerosis, where GWAS SNPs were more highly enriched among sQTLs than eQTLs (17). Furthermore, several studies have shown that sQTL analyses can uncover functions for GWAS-associated polymorphisms that would not have been identified through eQTL analysis alone. Zhang et al. found that 4.5% of GWAS SNPs from the GWAS catalog had evidence of cis-sQTLs but not cis-eQTLs (16). Li et al. showed that sQTLs identified in lymphoblastoid cell lines are enriched among autoimmune-disease associated variants (18). Another study demonstrated (17) that there were similar levels of enrichment of both eQTLs and sQTLs among SNPs associated with rheumatoid arthritis. In addition, Li et al. showed that in an analysis of rheumatoid arthritis, the inclusion of intronic splicing data allowed for the identification of 18 putative disease genes, of which 13 would not have been associated on the basis of gene-expression level measurements alone (17).

We identified seven top loci from GWAS that contained SNPs that colocalized with sQTLs. Of these loci, three could only be explained by sQTLs and not eQTLs. Of particular interest was the association at 5q32 in which SNPs in HTR4 were associated with COPD case-control status. This is a well-replicated GWAS association for lung function (33–35), COPD (11, 36) and airflow obstruction in smokers (10). Despite the consistent genetic association, as well as prior studies identifying *HTR4* expression in developing lung and increased airways resistance in a murine model, the mechanism by which the specific SNPs contribute to COPD risk is unknown, and additional genes in the region have not been investigated. Our analysis revealed that these SNPs may contribute to COPD by regulating splicing of a neighboring gene, *FBXO38*. This is consistent with evidence that the nearest gene to an associated SNP is the causative gene in only a minority of cases (37, 38). Therefore, analyses such as eQTL and sQTL studies are critical to uncover a relationship between an implicated SNP and the gene responsible for the association.

The characterization of splicing in F-box protein 38 (*FBXO38*) resulted in the discovery of a cryptic exon which has not been previously annotated. Through qPCR we were able to validate the existence of the exon as well as replicate the association with genotype in both blood and lung tissue. This is of particular importance as it demonstrates that analysis of splicing in the blood can uncover sQTLs that are also relevant to the lung. Sequence analysis using ORF Finder (39) determined that this newly identified transcript encodes a premature stop codon in the cryptic exon. This will result in a truncated protein that could have altered structure and function. To date, relatively little is known about the biological processes as well as the downstream signaling pathways through which FBXO38 operates to possibly influence COPD. Furthermore, no known ubiquitin substrate has been identified for FBXO38, which makes FBXO38 one of the orphan F-box proteins. Here we identified Cullin-1 as a binding partner for FBXO38. Therefore, the biological role of FBXO38 may rely mostly on its biochemical feature as an E3 ligase, through the formation of a complex with Cullin-1 and Skp1, similar to its family members Skp2 and Fbw7 (40, 41). *FBXO38* is additionally known to co-activate the transcriptional regulator *Kruppel Like Factor 7 (KLF7)* (41, 42). Members of the *KLF* gene family regulate cell proliferation, differentiation and survival, and have been found to play a role in airway inflammation (43). In particular, *KLF7* is involved in regulating epithelial cell differentiation and epithelial-mesenchymal transition (44, 45), and possibly in airway remodeling (46).

Another gene of interest is *SULT1A2*, located at 16p11.2. In this locus, SNPs were associated with both gene expression and splicing of *SULT1A2*. Therefore, two lines of evidence implicated *SULT1A2* as the causative gene. *SULT1A2* is a sulfotransferase enzyme, which is a family of phase II liver enzymes that detoxify a variety of endogenous and xenobiotic compounds (47). *SULT1A2* can sulfonate hormones like estrogens and androgens, but of particular interest for COPD, sulfation through *SULT1A2* is a pathway for the metabolism of cigarette smoke compounds (48). The 16p11.2 region contains a large inversion spanning ~0.45 MB. This region encompasses the entire *SULT1A2* gene as well as the *CCDC101* gene in which the associated SNPs are located, and is near other genes such as *TUFM*. This region contains SNPs which are associated with both obesity and body mass index (BMI)(49, 50) asthma (51), and autoimmune diseases like diabetes and inflammatory bowel disease; the inversion allele itself has been shown to be protective against the joint occurrence of asthma and obesity(52). There are 24 polymorphisms that have been shown accurately tag the inversion (52). The rs4788084 SNP which was identified in our study to be associated with both splicing and gene expression is among these markers, and is in LD with the inversion (R^2^=0.982). However, since both the SNP and the associated splice sites are located within the inversion, it is unlikely that inversion alters or contributes to this regulation of splicing.

Finally, we identified the *CDK11A* gene as a novel COPD candidate gene in the 1p36.32 locus. Here, SNPs were associated with the occurrence of a single exon in the *CDK11A* gene. *CDK11A* encodes a member of the serine/threonine protein kinase family. This kinase can be cleaved by caspases and may play a role in cell apoptosis (53, 54), which is a key dysregulated pathway in COPD (55, 56). As this locus did not reach genome-wide significance in the larger GWAS, additional studies will be needed to confirm the phenotypic association.

Most polymorphisms that were associated with splicing were acting from a distance, and were not located within or close to the splice donor or acceptor site. The majority of sQTLs (88%) were located in intronic or intergenic regions, and only 0.01% percent were located within splice sites. Furthermore, out of the seven GWAS loci that were discovered to have sQTLs, the most likely causative SNP was located greater than 500bp from the splice site in all cases. The mechanism by which these SNPs act on splicing is likely complex, but may involve auxiliary splicing regulatory elements such as exonic splicing enhancers (ESEs), intronic splicing enhancers (ISEs), exonic splicing silencers (ESSs) and intronic splicing silencers (ISSs). These regulatory elements control splicing through the recruitment of trans-acting factors that interact with other regulatory factors or core spliceosome components (57, 58).

A potential limitation of this study is that our eQTLs and sQTLs were identified in whole blood samples. COPD is a respiratory disease, and the most relevant cell-types in which to study gene expression changes may be located in the lung. However, due to the strong inflammatory component of COPD, characterization of gene expression and splicing in immune cells also has biological relevance. In addition, there is a high proportion of sQTL sharing across tissues (59), which has been recently supported by the finding that 75-93% of sQTLs are replicated across tissue pairs from the GTEx consortium, with the estimated level of sharing between whole blood and lung being 92% (17). Therefore, it is likely that the majority of the sQTLs identified in peripheral blood are also sQTLs in lung tissue. Furthermore, we experimentally demonstrated that the association with *FBXO38* splicing is also present in lung tissue, and therefore the mechanism discovered in whole blood is likely to also be of relevance in the lung. Another potential limitation is that peripheral blood samples contain a mixture of cell types, any of which could contribute to gene expression signals. We included white blood cell percentages as a covariate in eQTL and sQTL analyses to limit potential confounding by differences in cell proportions. An additional limitation of this study is the depth of sequencing of approximately 20 million reads per sample. This read depth was selected to maximize sequencing value within a large genetic epidemiology study. In order to be able capture the effect of genotype on gene expression and splicing, a large sample size was required, and therefore there was a tradeoff to be made between sequencing depth and sample size. While this could lead to the failure to capture rare splicing events, the goal of the study was to capture common splicing variations associated with genotype, which this design allowed us to do. Furthermore, it is important to note that the eCAVIAR calculation of CLPP is dependent on sample size as well as effect size of the eQTL/sQTL. As previously shown (30), an increase in sample size, or an increase in effect size results in higher CLPP values, and thus there is more power to detect colocalization with a larger sample size, or with eQTLs/sQTLs of strong effect. This means that while we may have been able to detect additional colocalization with a larger sample size, this does not impact our confidence in the CLPP values we have calculated here.

In conclusion, we found that many SNPs were associated with alternative splicing in peripheral blood. More COPD-associated variants were sQTLs than eQTLs, and we identified variant associations with splice sites in three genes including *FBXO38*, an orphan F-box protein, which have a function role in COPD through an effect as an E3 ligase on a currently unknown substrate. These data indicate that analysis of alternative splicing may provide novel insights into disease mechanisms.

## Supporting information

Supplementary Table 4: Results of eCaviar colocalization analysis between sQTLs/GWAS and eQTLs/GWAS for the seven GWAS loci containing sQTLs

## Acknowledgements

The authors thank the International COPD Genetics Consortium (ICGC) investigators for providing the genetic association data. We would also like to acknowledge the contributions of all COPDGene investigators. A complete list of ICGC and COPDGene investigators is available in the supplementary materials.

## Supporting Information Legends

### Supplementary Data File (word document)

Supplementary Table 1: Clinical characteristics of COPDGene study individuals included in the analysis.

Supplementary Table 2: cis eQTLs and sQTLs identified at the 10% FDR

Supplementary Table 3: Number of introns with start and stop sites that are annotated vs. cryptic.

Supplementary Table 5: KEGG pathways that are enriched in genes regulated by sQTLs but not eQTLs at the 5% FDR

Supplementary Table 6: Reactome pathways that are enriched in genes regulated by sQTLs but not eQTLs at the 5% FDR

Supplementary Figure 1: Schematic diagram illustrating the generation of splice clusters and calculation of splice ratios for 5q32

Supplementary Figure 2: Enrichment of low P-value associations with COPD case control status among sQTL and eQTL SNPs at the 10% FDR

Supplementary Figure 3: rs7730971 is associated with a cryptic splice site in FBXO38 Supplementary Figure 4: FBXO38 interacts with Cullin 1, but not other Cullin family members in 293T cells.

Supplementary Figure 5: rs4788084 is associated with a splice site in SULT1A2 as well as SULT1A2 whole gene expression

Supplementary Figure 6: rs2843126 is associated with a splice site in CDK11A

### Supplementary Table 4 (excel file)

Supplementary Table 1: Results of eCaviar colocalization analysis between sQTLs/GWAS and eQTLs/GWAS for the seven GWAS loci containing sQTLs

Author contributions
Conceptualization: AS, CPH; Data Curation: MHB, KdJ, THB, XZ, EKS, MHC, PJC, CPH; Formal Analysis: AS, JHY, MMP, RPC, AL, BDH; Writing – Original Draft Preparation: AS, CPH; Writing – Review & Editing: AS, JHY, MMP, RPC, AL, BDH, MHB, KdJ, THB, XZ, EKS, MHC, PJC, CPH

