## Supplementary Data File for "Analysis of genetically driven alternative splicing identifies FBXO38 as a novel COPD susceptibility gene"

**Supplementary Table 1: Clinical characteristics of COPDGene study individuals included in the analysis.**

|  | **Overall (n=376)** |
| --- | --- |
| Gender (% male) | 53.2 |
| Age, mean (SD) | 67.2 (8.4) |
| Current Smokers (% smokers) | 34.8 |
| Pack-Years Smoked, mean (SD) | 47.3 (23.5) |
| FEV1 percent predicted, mean (SD) | 73.6 (27.2) |
| Control - GOLD 0 (%)^1^ | 40.4 |
| COPD - GOLD 1 (%) | 9.8 |
| COPD - GOLD 2 (%) | 24.5 |
| COPD - GOLD 3 (%) | 16.2 |
| COPD - GOLD 4 (%) | 8.2 |
| Neutrophil percentage, mean (SD) | 61.8 (9.5) |
| Lymphocyte percentage, mean (SD) | 26.8 (8.6) |
| Eosinophil percentage, mean (SD) | 2.6 (2.2) |
| Monocyte percentage, mean (SD) | 8.2 (2.4) |

^1^Three individuals lack GOLD stage status due to lack of pulmonary function testing measurements

**Supplementary Table 2: cis eQTLs and sQTLs identified at the 10% FDR**

|  | **Cis-eQTL analysis** | **Cis-sQTL analysis** |
| --- | --- | --- |
| Tests conducted | 99,035,498 | 365,901,394 |
| Significant SNP-gene/intron pairs | 1,242,993 | 1,706,704 |
| Significant unique eQTL/sQTL SNPS | 708,928 | 561,060 |
| Significant genes/exons | 15,913 genes | 30,333 exons / 6742 genes |

**Supplementary Table 3: Number of introns with start and stop sites that are annotated vs. cryptic.**

| Verdict | Annotated to Genes^1^ |
| --- | --- |
| Fully Annotated | 18,951 |
| Cryptic 5’ splice site | 3,795 |
| Cryptic 3’ splice site | 3,986 |
| Cryptic Unanchored | 1,771 |
| Novel Annotated pair | 1,830 |

^1^Introns for which a most likely gene could be identified based on intron start and stop sites.

**Supplementary Table 5: KEGG pathways that are enriched in genes regulated by sQTLs but not eQTLs at the 5% FDR**

| Pathway | Number of genes supporting pathway | P-value | Bonferroni adjusted P-value |
| --- | --- | --- | --- |
| RNA transport | 31 | 9.49E-143 | 2.66E-140 |
| Endocytosis | 120 | 1.48E-75 | 4.16E-73 |
| Ubiquitin mediated proteolysis | 21 | 4.81E-63 | 1.35E-60 |
| Protein processing in endoplasmic reticulum | 99 | 2.59E-39 | 7.26E-37 |
| Oocyte meiosis | 136 | 8.89E-22 | 2.49E-19 |
| Prostate cancer | 180 | 2.29E-19 | 6.42E-17 |
| Lysosome | 17 | 1.43E-16 | 4.01E-14 |
| Peroxisome | 22 | 5.37E-13 | 1.50E-10 |
| Aminoacyl-tRNA biosynthesis | 10 | 9.02E-13 | 2.53E-10 |
| Prolactin signaling pathway | 160 | 3.03E-11 | 8.47E-09 |
| Renal cell carcinoma | 173 | 3.10E-11 | 8.67E-09 |
| Oxidative phosphorylation | 64 | 6.75E-11 | 1.89E-08 |
| mRNA surveillance pathway | 81 | 1.25E-10 | 3.49E-08 |
| Alzheimer’s disease | 60 | 3.43E-09 | 9.60E-07 |
| Ribosome biogenesis in eukaryotes | 78 | 4.39E-08 | 1.23E-05 |
| NOD-like receptor signaling pathway | 106 | 1.80E-07 | 5.03E-05 |
| RNA degradation | 12 | 4.72E-07 | 0.000132 |
| Fc gamma R-mediated phagocytosis | 96 | 5.80E-07 | 0.0001623 |
| Colorectal cancer | 129 | 6.39E-07 | 0.0001788 |
| SNARE interactions in vesicular transport | 15 | 9.30E-07 | 0.0002603 |

**Supplementary Table 6: Reactome pathways that are enriched in genes regulated by sQTLs but not eQTLs at the 5% FDR**

| Pathway | Number of genes supporting pathway | P-value | Bonferroni adjusted P-value |
| --- | --- | --- | --- |
| Golgi to ER Retrograde Transport | 53 | 1.42E-24 | 1.25E-21 |
| Membrane Trafficking | 32 | 4.70E-21 | 4.14E-18 |
| PPARA activates gene expression | 17 | 7.39E-18 | 6.51E-15 |
| mRNA Splicing | 15 | 3.38E-17 | 2.97E-14 |
| Respiratory electron transport | 17 | 3.38E-16 | 2.97E-13 |
| SCF-beta-TrCP mediated degradation of Emi1 | 77 | 4.29E-15 | 3.77E-12 |
| Rev-mediated nuclear export of HIV RNA | 30 | 5.60E-15 | 4.93E-12 |
| RAF activation | 91 | 5.35E-13 | 4.70E-10 |
| Synthesis of substrates in N-glycan biosynthesis | 30 | 6.79E-13 | 5.97E-10 |
| Viral Messenger RNA Synthesis | 30 | 2.87E-12 | 2.52E-09 |
| Transcription of the HIV genome | 50 | 1.10E-11 | 9.64E-09 |
| Downstream signaling of activated FGFR1 | 35 | 7.77E-11 | 6.84E-08 |
| Organelle biogenesis and maintenance | 48 | 8.46E-11 | 7.45E-08 |
| Signaling by SCF-KIT | 30 | 1.61E-10 | 1.42E-07 |
| Asparagine N-linked glycosylation | 18 | 2.56E-10 | 2.25E-07 |
| Downstream signaling of activated FGFR4 | 59 | 3.52E-10 | 3.09E-07 |
| PI-3K cascade:FGFR2 | 18 | 3.63E-10 | 3.19E-07 |
| Initiation of Nuclear Envelope Reformation | 26 | 4.71E-10 | 4.15E-07 |
| Infectious disease | 189 | 5.47E-10 | 4.81E-07 |
| trans-Golgi Network Vesicle Budding | 16 | 6.32E-10 | 5.56E-07 |
| Endosomal Sorting Complex Required For Transport (ESCRT) | 22 | 6.77E-10 | 5.96E-07 |
| Post-translational protein modification | 49 | 7.41E-10 | 6.52E-07 |
| Transport to the Golgi and subsequent modification | 8 | 1.25E-09 | 1.10E-06 |
| KSRP (KHSRP) binds and destabilizes mRNA | 69 | 1.29E-09 | 1.13E-06 |
| Golgi Associated Vesicle Biogenesis | 11 | 1.43E-09 | 1.26E-06 |
| Toll Like Receptor 9 (TLR9) Cascade | 40 | 1.99E-09 | 1.75E-06 |
| Respiratory electron transport, ATP synthesis by chemiosmotic coupling, and heat production by uncoupling proteins. | 21 | 2.24E-09 | 1.97E-06 |
| BMAL1:CLOCK,NPAS2 activates circadian gene expression | 61 | 1.31E-08 | 1.15E-05 |
| Lysosome Vesicle Biogenesis | 9 | 7.77E-08 | 6.84E-05 |
| Toll Like Receptor TLR1:TLR2 Cascade | 34 | 8.46E-08 | 7.44E-05 |
| APC:Cdc20 mediated degradation of cell cycle proteins prior to satisfation of the cell cycle checkpoint | 31 | 9.07E-08 | 7.98E-05 |
| Sema4D in semaphorin signaling | 61 | 9.40E-08 | 8.27E-05 |
| SUMOylation | 13 | 1.12E-07 | 9.82E-05 |
| S45 mutants of beta-catenin aren’t phosphorylated | 33 | 1.45E-07 | 0.0001275 |
| Antigen processing: Ubiquitination & Proteasome degradation | 20 | 3.93E-07 | 0.0003459 |

**Supplementary Figure 1: Schematic diagram illustrating the generation of splice clusters and calculation of splice ratios for 5q32**

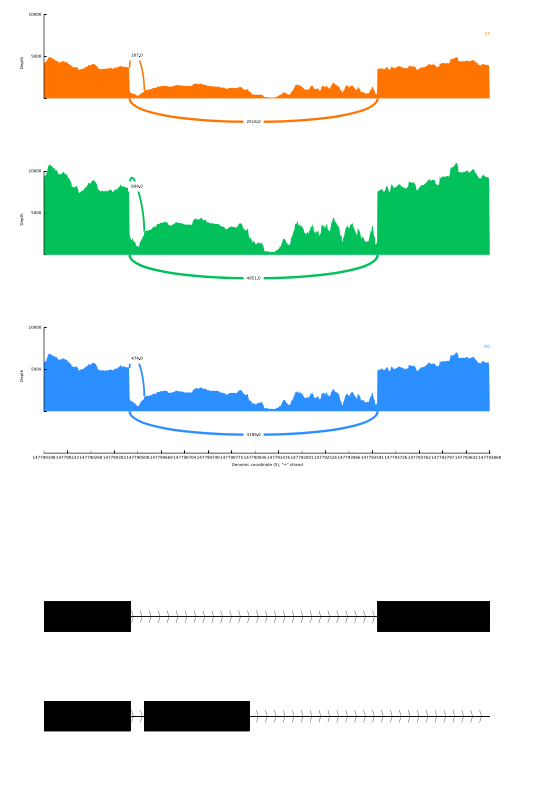

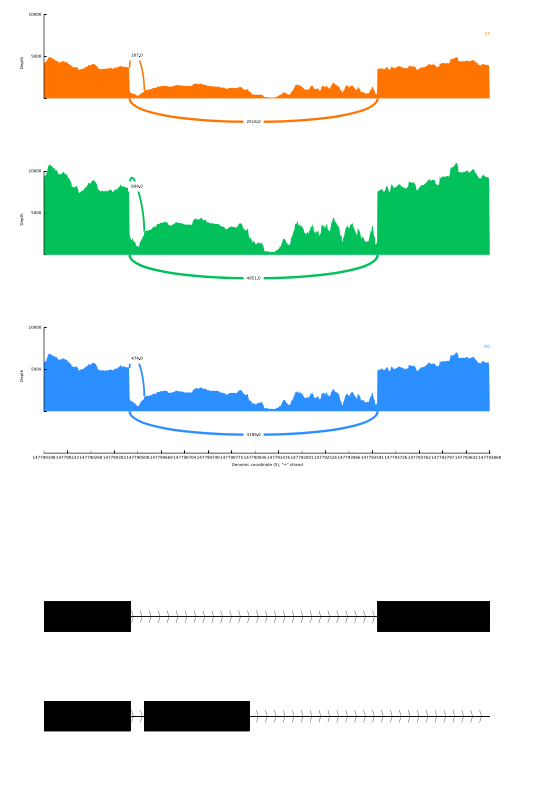

Splice site 1

Splice site 3

Splice site 2

Exon 9

Exon 10

Exon 9

Exon 10

Novel Exon

FBXO38 Isoforms

RNASeq reads mapped to genome

Cluster reads according to shared intron start and stop positions

Cluster clu_10408

Splice site 1

Calculate splice ratios

Splice site 2

Splice site 3

Splice site 1

Splice site 1 + 2 + 3

Splice site 2

Splice site 1 + 2 + 3

Splice site 3

Splice site 1 + 2 + 3

RNA-seq reads are mapped to the genome, and reads align to both previously annotated as well as undocumented exons. Using leafcutter, reads are clustered together based on shared start and stop intron positions. Splice ratios are then calculated by dividing the number of reads that support the presence of a given junction by the total number of reads in the cluster. This ratio is used as the input for

**Supplementary Figure 2: Enrichment of low P-value associations with COPD case control status among sQTL and eQTL SNPs at the 10% FDR**

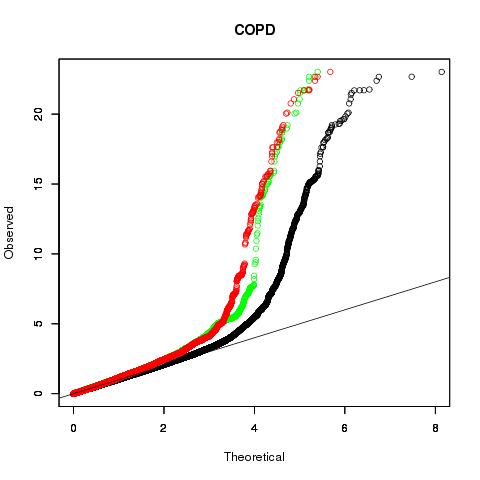

sQTLs (558,660)

eQTLs (706,030)

Genome-wide

COPD Case/Control GWAS data

**Supplementary Figure 3: rs7730971 is associated with a cryptic splice site in FBXO38**

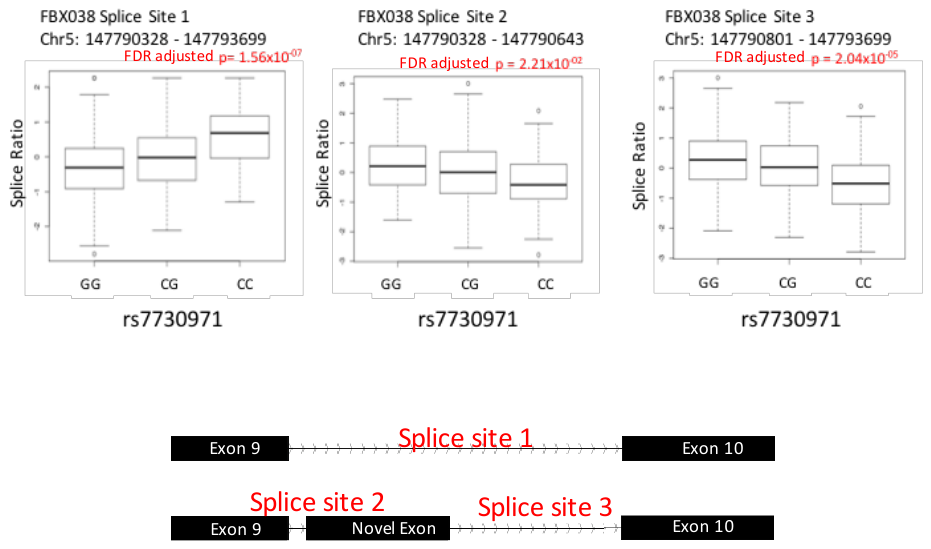

**Supplementary Figure 4: FBXO38 interacts with Cullin 1, but not other Cullin family members in 293T cells.**

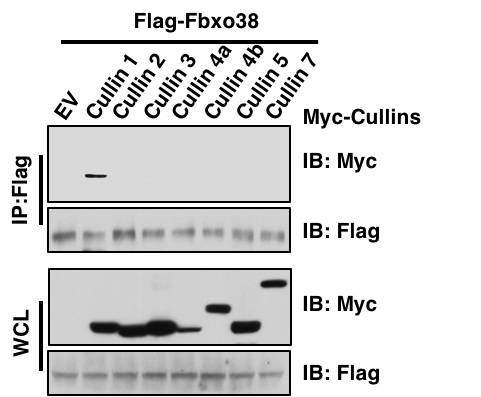

**Supplementary Figure 5: rs4788084 is associated with a splice site in SULT1A2 as well as SULT1A2 whole gene expression**

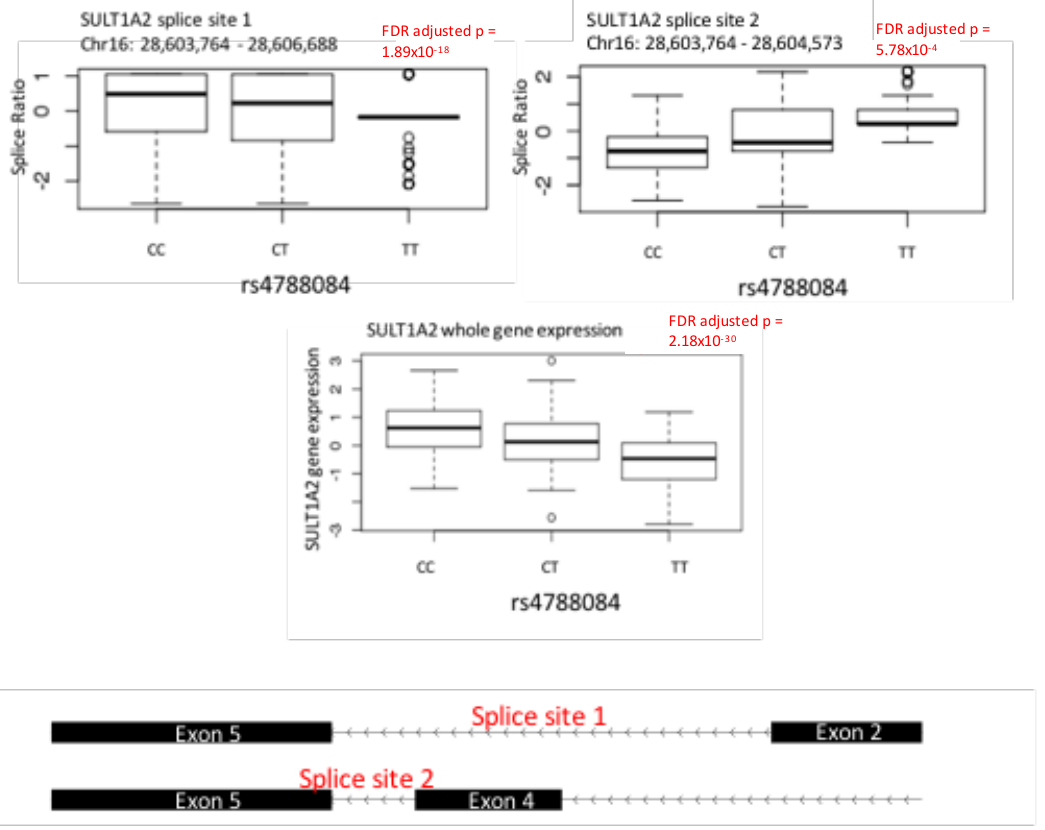

**Supplementary Figure 6: rs2843126 is associated with a splice site in CDK11A**

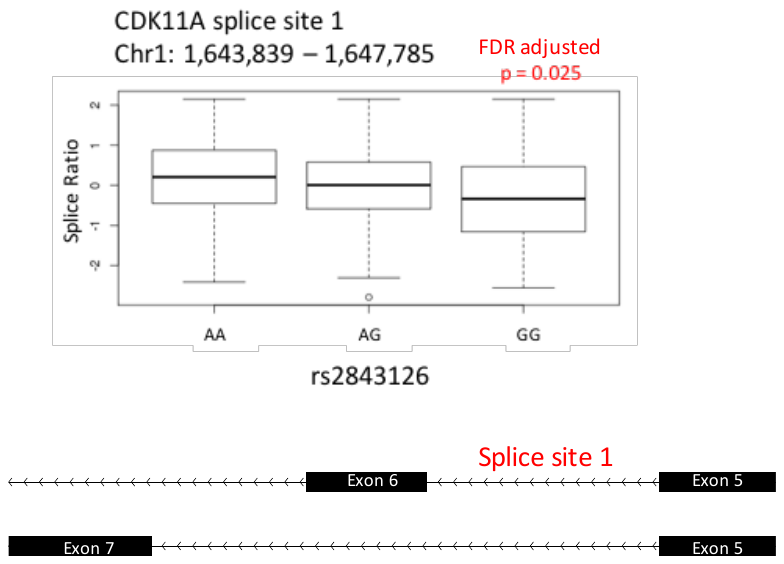

**Acknowledgements:**

**COPDGene Investigators – Core Units**

*Administrative Center*: James D. Crapo, MD (PI); Edwin K. Silverman, MD, PhD (PI); Barry J. Make, MD; Elizabeth A. Regan, MD, PhD

*Genetic Analysis Center*: Terri Beaty, PhD; Ferdouse Begum, PhD; Robert Busch, MD; Peter J. Castaldi, MD, MSc; Michael Cho, MD; Dawn L. DeMeo, MD, MPH; Adel R. Boueiz, MD; Marilyn G. Foreman, MD, MS; Eitan Halper-Stromberg; Nadia N. Hansel, MD, MPH; Megan E. Hardin, MD; Lystra P. Hayden, MD, MMSc; Craig P. Hersh, MD, MPH; Jacqueline Hetmanski, MS, MPH; Brian D. Hobbs, MD; John E. Hokanson, MPH, PhD; Nan Laird, PhD; Christoph Lange, PhD; Sharon M. Lutz, PhD; Merry-Lynn McDonald, PhD; Margaret M. Parker, PhD; Dandi Qiao, PhD; Elizabeth A. Regan, MD, PhD; Stephanie Santorico, PhD; Edwin K. Silverman, MD, PhD; Emily S. Wan, MD; Sungho Won

*Imaging Center*: Mustafa Al Qaisi, MD; Harvey O. Coxson, PhD; Teresa Gray; MeiLan K. Han, MD, MS; Eric A. Hoffman, PhD; Stephen Humphries, PhD; Francine L. Jacobson, MD, MPH; Philip F. Judy, PhD; Ella A. Kazerooni, MD; Alex Kluiber; David A. Lynch, MB; John D. Newell, Jr., MD; Elizabeth A. Regan, MD, PhD; James C. Ross, PhD; Raul San Jose Estepar, PhD; Joyce Schroeder, MD; Jered Sieren; Douglas Stinson; Berend C. Stoel, PhD; Juerg Tschirren, PhD; Edwin Van Beek, MD, PhD; Bram van Ginneken, PhD; Eva van Rikxoort, PhD; George Washko, MD; Carla G. Wilson, MS;

*PFT QA Center, Salt Lake City, UT*: Robert Jensen, PhD

*Data Coordinating Center and Biostatistics*, *National Jewish Health, Denver, CO*: Douglas Everett, PhD; Jim Crooks, PhD; Camille Moore, PhD; Matt Strand, PhD; Carla G. Wilson, MS

*Epidemiology Core*, *University of Colorado Anschutz Medical Campus, Aurora, CO*: John E. Hokanson, MPH, PhD; John Hughes, PhD; Gregory Kinney, MPH, PhD; Sharon M. Lutz, PhD; Katherine Pratte, MSPH; Kendra A. Young, PhD

**COPDGene Investigators – Clinical Centers**

*Ann Arbor VA:* Jeffrey L. Curtis, MD; Carlos H. Martinez, MD, MPH; Perry G. Pernicano, MD

*Baylor College of Medicine, Houston, TX*: Nicola Hanania, MD, MS; Philip Alapat, MD; Mustafa Atik, MD; Venkata Bandi, MD; Aladin Boriek, PhD; Kalpatha Guntupalli, MD; Elizabeth Guy, MD; Arun Nachiappan, MD; Amit Parulekar, MD;

*Brigham and Women’s Hospital, Boston, MA*: Dawn L. DeMeo, MD, MPH; Craig Hersh, MD, MPH; Francine L. Jacobson, MD, MPH; George Washko, MD

*Columbia University, New York, NY*: R. Graham Barr, MD, DrPH; John Austin, MD; Belinda D’Souza, MD; Gregory D.N. Pearson, MD; Anna Rozenshtein, MD, MPH, FACR; Byron Thomashow, MD

*Duke University Medical Center, Durham, NC*: Neil MacIntyre, Jr., MD; H. Page McAdams, MD; Lacey Washington, MD

*HealthPartners Research Institute, Minneapolis, MN*: Charlene McEvoy, MD, MPH; Joseph Tashjian, MD

*Johns Hopkins University, Baltimore, MD*: Robert Wise, MD; Robert Brown, MD; Nadia N. Hansel, MD, MPH; Karen Horton, MD; Allison Lambert, MD, MHS; Nirupama Putcha, MD, MHS

*Los Angeles Biomedical Research Institute at Harbor UCLA Medical Center, Torrance, CA*: Richard Casaburi, PhD, MD; Alessandra Adami, PhD; Matthew Budoff, MD; Hans Fischer, MD; Janos Porszasz, MD, PhD; Harry Rossiter, PhD; William Stringer, MD

*Michael E. DeBakey VAMC, Houston*, *TX*: Amir Sharafkhaneh, MD, PhD; Charlie Lan, DO

*Minneapolis VA:* Christine Wendt, MD; Brian Bell, MD

*Morehouse School of Medicine, Atlanta, GA*: Marilyn G. Foreman, MD, MS; Eugene Berkowitz, MD, PhD; Gloria Westney, MD, MS

*National Jewish Health, Denver, CO*: Russell Bowler, MD, PhD; David A. Lynch, MB

*Reliant Medical Group, Worcester, MA*: Richard Rosiello, MD; David Pace, MD

*Temple University, Philadelphia, PA:* Gerard Criner, MD; David Ciccolella, MD; Francis Cordova, MD; Chandra Dass, MD; Gilbert D’Alonzo, DO; Parag Desai, MD; Michael Jacobs, PharmD; Steven Kelsen, MD, PhD; Victor Kim, MD; A. James Mamary, MD; Nathaniel Marchetti, DO; Aditi Satti, MD; Kartik Shenoy, MD; Robert M. Steiner, MD; Alex Swift, MD; Irene Swift, MD; Maria Elena Vega-Sanchez, MD

*University of Alabama, Birmingham, AL:* Mark Dransfield, MD; William Bailey, MD; Surya Bhatt, MD; Anand Iyer, MD; Hrudaya Nath, MD; J. Michael Wells, MD

*University of California, San Diego, CA*: Joe Ramsdell, MD; Paul Friedman, MD; Xavier Soler, MD, PhD; Andrew Yen, MD

*University of Iowa, Iowa City, IA*: Alejandro P. Comellas, MD; John Newell, Jr., MD; Brad Thompson, MD

*University of Michigan, Ann Arbor, MI*: MeiLan K. Han, MD, MS; Ella Kazerooni, MD; Carlos H. Martinez, MD, MPH

*University of Minnesota, Minneapolis, MN*: Joanne Billings, MD; Abbie Begnaud, MD; Tadashi Allen, MD

*University of Pittsburgh, Pittsburgh, PA*: Frank Sciurba, MD; Jessica Bon, MD; Divay Chandra, MD, MSc; Carl Fuhrman, MD; Joel Weissfeld, MD, MPH

*University of Texas Health Science Center at San Antonio, San Antonio, TX*: Antonio Anzueto, MD; Sandra Adams, MD; Diego Maselli-Caceres, MD; Mario E. Ruiz, MD

**ICGC Executive Committee**: James Crapo, William MacNee, David Lynch, Dirkje Postma, Edwin Silverman, Jorgen Vestbo. *Members*: Alvar Agusti, Wayne Anderson, Nawar Bakerly, Per Bakke, Robert Bals, Kathleen Barnes, R. Graham Barr, Terri Beaty, Eugene Bleecker, Marike Boezen, Yohan Bosse, Russell Bowler, Christopher Brightling, Marleen de Bruijne, Guy Brusselle, Peter Castaldi, Bartolome Celli, Michael Cho, Harvey Coxson, Ron Crystal, Dawn DeMeo, Asger Dirksen, Jennifer Dy, Marilyn Foreman, Judith Garcia-Aymerich, Pierre Gevenois, Hester Gietema, Ian Hall, Nadia Hansel, Craig Hersh, Eric Hoffman, John Hokanson, Pim de Jong, Phil Judy, Noor Kalsheker, Hans-Ulrich Kauczor, Woo Jin Kim, Lies Lahousse, Tarja Laitinen, Diether Lambrechts, Sang Do Lee, Augusto Litonjua, David Lomas, Stephanie London, Daan Loth, Sharon Lutz, Masaharu Nishimura, John Newell, Merry-Lynn McDonald, Borge Nordestgaard, George O'Connor, Yeon-Mok Oh, Peter Pare, Massimo Pistolesi, Milo Puhan, Elizabeth Regan, Stephen Rennard, Stephen Rich, Andrew Sandford, Joon-Beom Seo, Andrea Short, Martijn Spruit, Berend Stoel, David Strachan, Gerben ter Riet, Nicola Sverzellati, Yohannes Tesfaigzi, Martin Tobin, Edwin Van Beek, Bram van Ginneken, Claus Vogelmeier, Adam Wanner, George Washko, Els Wauters, Emiel Wouters, Robert Young, and Loems Zeigler-Heitbrock
